# A Scalable Framework for Pan-Cancer Tumor Evolution Analysis Enables Transfer of Progression Mechanisms Across Tumor Entities

**DOI:** 10.64898/2026.01.20.700556

**Authors:** Simon Pfahler, Andreas Lösch, Y. Linda Hu, Rudolf Schill, Rainer Spang, Tilo Wettig

## Abstract

**Motivation:** Cancer progression is driven by the stochastic accumulation of interacting genetic alterations, and tumors from different tissues can share common evolutionary patterns despite distinct anatomical origins. While simplified progression models can be applied at scale, the use of expressive models that capture complex inter-event dependencies remains challenging in pan-cancer settings due to their computational demands.

**Results:** We present fastMHN, a scalable approximation to Mutual Hazard Network-based cancer progression models, that enables pan-cancer analysis of large genomic datasets with the explicit aim of enabling transfer of progression mechanisms across tumor entities. Applying fastMHN to a large clinical cohort, we identify a tumor-progression group spanning multiple tissues that is characterized by a link between STK11 mutations and poor patient survival. While the clinical relevance of STK11 mutations is well established in non-small cell lung cancer, our results suggest that a similar progression mechanism is present in molecularly defined subgroups of other cancer types. These findings illustrate how scalable pan-cancer progression modeling can facilitate cross-entity transfer of biological and potentially clinical insights.

**Availability and implementation:** The pan-cancer classification workflow and all data are available at github.com/simon-pfahler/fastMHN-classification.

## INTRODUCTION

### Motivation

Cancer is traditionally classified by tissue or organ of origin, such as lung, breast, colorectal, or prostate cancer, and treatment recommendations are largely organ specific [47]. While this classification remains clinically central, it does not fully reflect the molecular patterns observed within and across histopathological cancer types [21, 47]. At the genomic level, tumors arising from the same tissue can differ substantially, whereas tumors from different tissues can share highly similar genetic alterations and pathways [22]. This observation has motivated molecular and pan-cancer classifications that group tumors by shared genetic features rather than anatomical origin alone [27].

Tumor genetics is not static but evolves over time through a stochastic process in which somatic mutations and copy-number alterations accumulate and interact [19]. Cancer progression models aim to capture these dynamics explicitly, describing how genetic events influence the probability of subsequent alterations [12]. While individual tumors follow distinct evolutionary trajectories, similarities in these dynamics can be found across tumors of different tissue origin [27]. Mutual Hazard Networks (MHN) [40] and their extensions, such as observation Mutual Hazard Networks (oMHN) [39], represent state-of-the-art probabilistic models of cancer progression. They encode how the occurrence of one genetic event modulates the hazard rates of others, allowing for both promoting and inhibiting interactions. Applied in a pan-cancer setting, such models have the potential to uncover shared progression pathways that are not apparent in analyses based on cancer types. However, direct application of MHN-based models to pan-cancer datasets is computationally infeasible. Exact model training scales exponentially with the number of progression events and linearly with the number of samples [40], prohibiting analysis of datasets with many genes and thousands of tumors.

The resulting computational challenge is to develop approximation methods that preserve the expressive power of MHN-based progression models while making pan-cancer training tractable. Here, we exploit the common assumption that cancer progression is weakly modular, i.e., that genetic events form clusters with strong internal interactions, with only weak interactions between clusters [8, 44].

Building on this assumption, we introduce fastMHN, a scalable approximation to oMHN training that decomposes tumor-specific genetic events into weakly interacting clusters. This reduces computational complexity drastically while keeping approximation errors controlled, enabling MHN-based pan-cancer analysis (here, specifically solid cancers) for the first time. Applying fastMHN to the MSK-CHORD dataset [23] of more than ten thousand tumors across five tissues of origin, we uncover shared progression dynamics that recur across molecularly defined subgroups of different tumor types.

In particular, our analysis highlights a progression mechanism characterized by STK11 mutations that is associated with reduced patient survival. While this mechanism is well established in non-small cell lung cancer [37], our results suggest that it also operates in subsets of other tumor entities, where it has so far remained largely unexplored. More generally, these findings illustrate how pan-cancer tumor evolution analysis can support systematic transfer of biological and clinical knowledge across oncological fields, enabling insights from one cancer type to inform interpretation and decision making in others.

### Related work

Approximate training of MHNs has been studied since the formulation of the model itself. Most prominently, low-rank tensor networks [20] were used with some success to reduce the computational complexity from exponential to polynomial in the number of events [17, 36], but have never been successfully applied to biological data. Our strategy of utilizing weak modularity to obtain approximations is closely related to the junction-tree algorithm [1, 30], which is exact for stationary probability distributions but breaks down when introducing a time dependency. The method we present is built upon studies of the Sylvester equation, in particular the ADI iteration [4].

De-novo classification of pan-cancer tumor samples has been studied extensively in the literature. Foundational work was done in [22, 41], where latent-space methods were used to classify samples based on multi-omics data. There are classification methods based on various off-the-shelf statistical and machine-learning methods, using various types of multi-omics data [9, 25, 28, 46, 50]. FastMHN is most closely related to the classification method proposed in [27], which uses conjunctive Bayesian networks (CBN) [3] to learn progression dynamics from mutational data.

## METHODS

We start this section with a brief introduction to MHN [40] and an extension, oMHN [39]. We then discuss the strategy we employ to keep the computational cost manageable even when many genetic events are active. At the end of the section, we explain how we use this strategy to cluster pan-cancer samples into de-novo groups based on oMHNs.

### Mutual Hazard Network

Cancer progression is governed by the accumulation of genetic events such as mutations or copy-number alterations [32]. In this work we consider only binary events, where the state of an event is represented by 0 (absent) or 1 (present). A tumor with *d* possible events can thus be represented by a *d*-dimensional vector *x* ∈ {0, 1} ^*d*^. Using MHN, promoting and inhibiting influences between pairs of events can be inferred. This training is done on a dataset 𝒟 of tumors observed in patients.

MHN models the process of cancer progression as a continuous-time Markov chain, with the assumptions that no events are active at time *t* = 0, that events occur only one at a time, and that events are irreversible. The rates for the transitions from 0 to 1 are given by the proportional hazard assumption [10], resulting in a transition-rate matrix

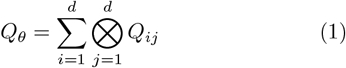

with

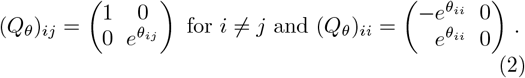

The *Q* matrix depends on a parameter matrix *θ* ∈ ℝ^*d×d*^, where 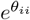 is the base rate of event *i* occurring, and 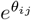 is the multiplicative influence of event *j* on event *i*. MHN allows for promoting (*θ*_*ij*_ > 0) as well as inhibiting (*θ*_*ij*_ < 0) effects between pairs of events.

In Markov-chain theory the probability distribution at time *t* ≥ 0 is given by

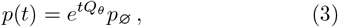

where 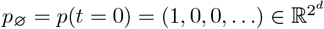.

The datasets we work with do not contain the tumor age *t*, which generally is an unknown quantity. Therefore we marginalize over *t*, assuming *t* to be an exponentially distributed random variable with mean 1,^1^ resulting in the time-marginalized probability distribution

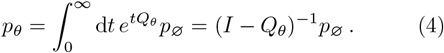

We can now fit a parameter matrix *θ* to best describe the data by maximizing the log-likelihood of observing the dataset 𝒟 under our model *θ* [5],

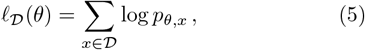

where *p*_*θ,x*_ is the entry of the vector *p*_*θ*_ at position *x* (with *x* mapped to 1, …, 2^*d*^). This optimization is usually done via algorithms such as BFGS [14] or Adam [26], which need not only the score 𝓁 _𝒟_ (*θ*) but also its gradient with respect to the parameters *θ*. Calculating score and gradient is computationally expensive, as exponentially large linear equations like Eq. (4) have to be solved [40]. We will address this problem below.

### Observation MHN

Most cancer progression models, including MHN, suffer from observational bias [18, 38, 40]. For MHN, an extension called observation MHN (oMHN) has been introduced in [39] to alleviate this bias. In oMHN, an additional event is introduced which represents the observation of the tumor and stops the progression. We are now interested in the stationary probability distribution *p*(∞). Ref. [39] arrives at a formula similar to Eq. (4),

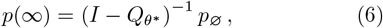

where

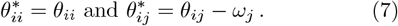

The additional parameters *ω*_*j*_ for *j* ∈ {1, …, *d*} denote the multiplicative influence of event *j* on the observation. Using oMHN instead of MHN is beneficial since effects due to observation bias are now explained via the observation rates and no longer via pairwise influences.

### Approximate MHN training with clustering

The computational complexity of MHN typically restricts us to *d*_max_ ≲25 active events per sample. For pan-cancer applications, this bound prevents us from applying the method to meaningful datasets. Therefore, we need approximations to construct MHNs for datasets that contain samples with more than *d*_max_ events.

To this end, we introduce a partition *P* = {*C*_1_, …, *C*_*n*_}, which is a set of clusters *C*_*k*_ ⊂ {1, …, *d*} such that every *i* ∈ {1, …, *d*} is in one and only one cluster *C*_*k*_. Now, consider a sample *x* with *d*_active_(*x*) > *d*_max_ active events and define

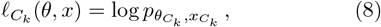

where 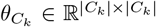 and 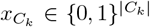 are the restrictions of *θ* and *x* to the cluster *C*_*k*_. We also define the sum of 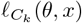 over all clusters of the partition *P* as

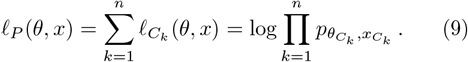

This yields an approximation

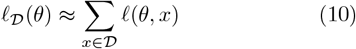

to the log-likelihood (5), where

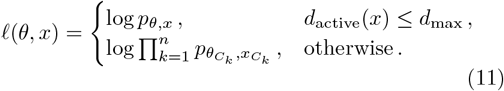

For samples with *d*_active_(*x*) > *d*_max_ this is the first-order approximation in the ADI iteration [4, 15, 29, 43]. Using this strategy we can now efficiently obtain approximations to the log-likelihood and gradient of the MHN in 𝒪 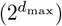 time.

To obtain a partition *P* for a sample *x* ∈ 𝒟 we perform a single-linkage hierarchical clustering of events, based on the current MHN. For the distance between two events we use

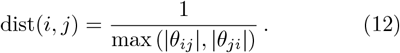

Whenever the sum of active events for sample *x* in two clusters is larger than *d*_max_, we explicitly set their distance to ∞ to make sure that no cluster contains more than *d*_max_ events.

The relative error of the approximations of the log-likelihood contributions in Eq. (10) and their gradients is shown in Fig. 1. We also compare the log-likelihood of MHNs trained via this strategy to results without the approximation, see Fig. 2. In both cases, the quality of the approximation increases with *d*_max_ as expected.

**Figure 1.**
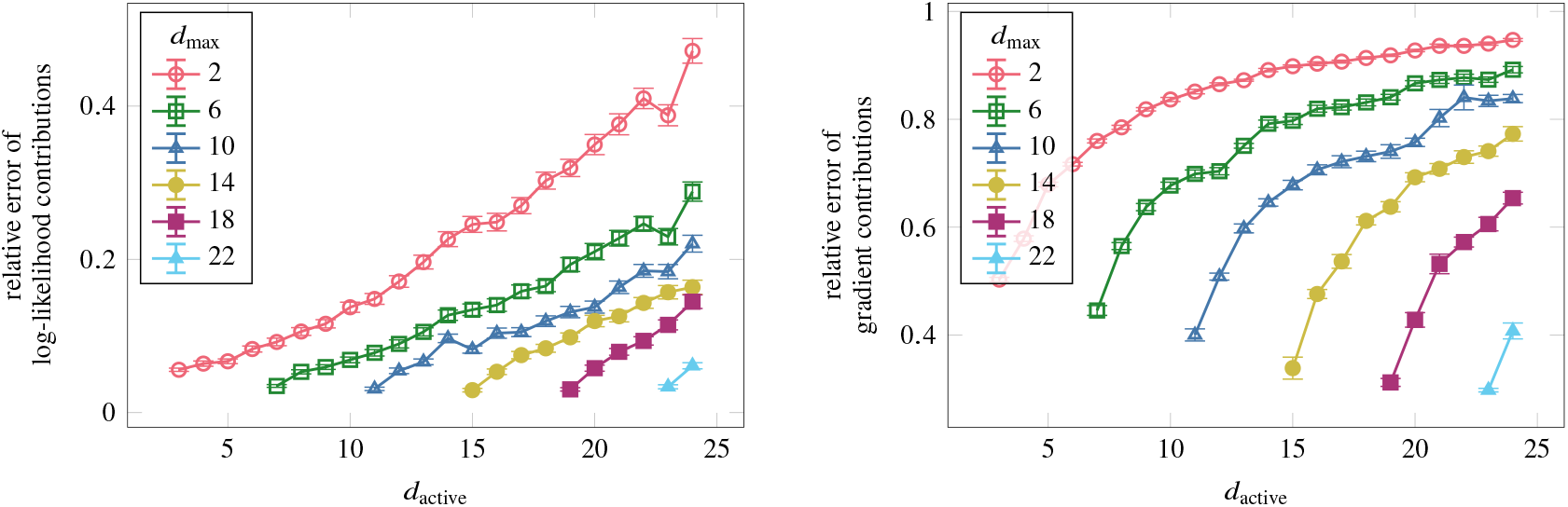
Quality of the approximations of the MHN log-likelihood (left) and its gradient (right), averaged over all contributions with fixed *d*_active_. Data are obtained from artificial datasets with a total of 256 events, but only samples with ≤ 24 active events are shown, since for larger samples no exact result is available. Only data points where the approximation is used are shown.

**Figure 2.**
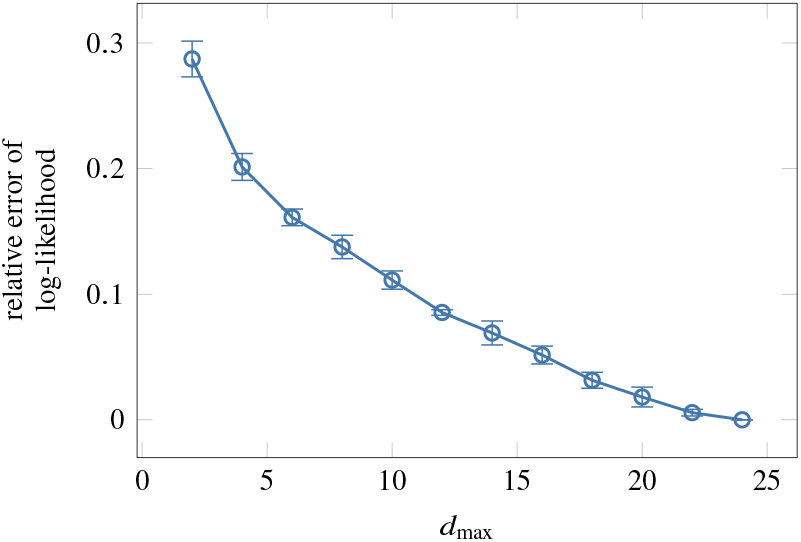
Quality of MHNs obtained via the approximate training strategy. Data are obtained as described in Fig. 1, again with *d*_active_ ≤ 24.

We tested various distance functions and found that Eq. (12) gave the best results in terms of quality of the approximations (see Fig. 1), quality of trained MHNs (see Fig. 2), and accuracy of pan-cancer classifications (see Fig. 3).

**Figure 3.**
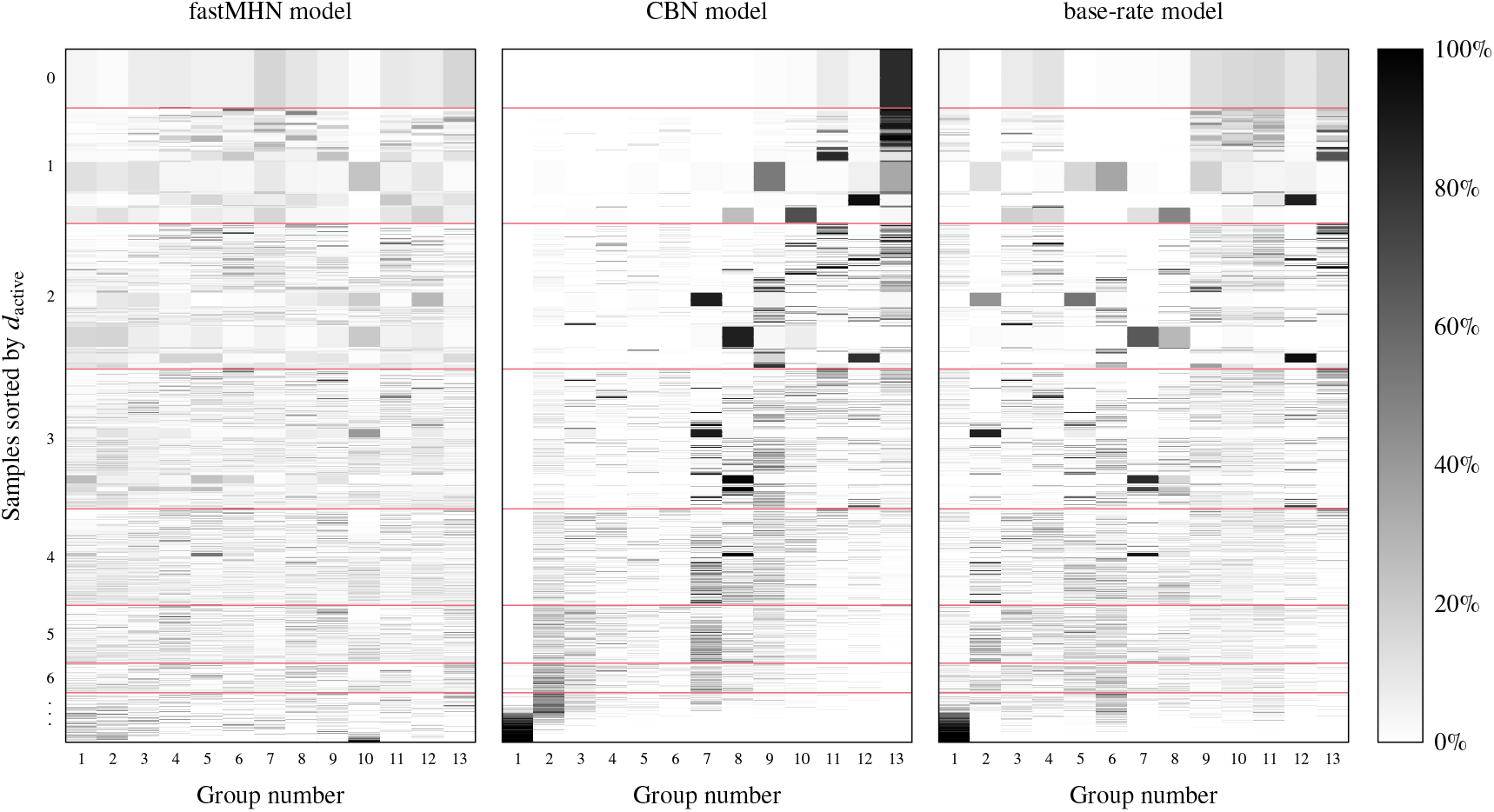
Membership probabilities for our fastMHN model, for the CBN model [27], and for the base-rate model. All models are shown for the same number of groups to simplify comparisons between the models. Groups are sorted by average *d*_active_ of the groups. Each row of the plot shows the membership probabilities of a single sample. The horizontal lines indicate where the number of active events changes.

### Pan-cancer classification

We now apply the fastMHN method of inferring approximate oMHNs to the task of finding novel groups of cancer across different tissues. For classifying |𝒟| pan-cancer samples into *n* groups, we initialize the group membership probabilities 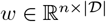 uniformly and add a small Gaussian noise to those probabilities. Then, an iterative expectation-maximization algorithm [31, 34] is performed, consisting of the following expectation (E) and maximization (M) steps:

E For each group *k*, an oMHN parameterized by *θ*_*k*_ is learned via 5-fold cross-validation [5] using all samples and this group’s membership probabilities *w*_*k,x*_. To include *w*_*k,x*_ in the log-likelihood of the oMHN of group *k*, every term in the sum in Eq. (10) is weighted by *w*_*k,x*_.

M The membership probabilities are updated according to the likelihoods of the samples under the learned oMHN models,

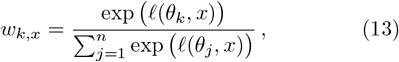

where 𝓁 (*θ, x*) is the term in the sum of Eq. (10).

Uniform group membership probabilities constitute an unstable equilibrium of this algorithm [34]. Our starting point is close to uniform, and hence the groups quickly start to differ from one another. The iterative process is repeated until the membership probabilities become stable again.

One such classification process includes the training of a large number of oMHNs.^2^ Therefore, oMHN without the approximation is limited to datasets with 15 active events per sample.

The quality of a classification is evaluated by calculating its Akaike information criterion [6], which for a dataset with *d* events is given by

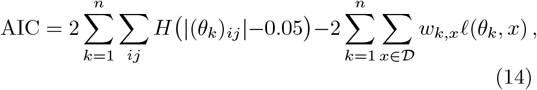

where *H*(*x*) is the Heaviside step function. The first term is the effective number of parameters of the model [35], and the second is its log-likelihood.

## RESULTS

The approximation strategy introduced above removes the key computational bottleneck that has previously prevented the application of oMHN to large pan-cancer datasets. As a result, it is now possible to investigate progression dynamics across thousands of tumors and dozens of active genetic events. In the following, we show which biological patterns become visible through this analysis.

### Data preparation

We apply the presented methods to find novel groups from pan-cancer samples in the MSK-CHORD dataset [23]. This dataset includes bulk sequencing data along with histopathological annotations. Via cbioportal [7, 11, 16], binary mutation data and histopathological annotations were accessed. We consider only primary tumor samples and only samples from patients who do not have primary tumor samples from multiple cancer types. The resulting dataset contains a total of 15 816 samples across the cancer types breast (2844 samples), colorectal (4027 samples), non-small cell lung (4866 samples), pancreatic (1964 samples), and prostate (2115 samples). We use the 15 most common genetic mutations (grouped by gene) for each of the 5 cancer types, resulting in a gene panel of 47 events total. Note that by simply mentioning a gene name we imply the event of a mutation in that gene. The maximum *d*_active_ in the dataset is 38, which is far beyond the previous limitation of oMHN [45].

### Statistical model quality

We perform classification into a varying number *n* of groups using fastMHN. For comparison, we also perform classification using the CBN method described in [2, 27]. As a further baseline, we also consider an MHN model that does not allow for any promoting or inhibiting effects, modeling all events as independent. We call this the base-rate model, as it is an MHN model that only contains base rates.

For fastMHN training processes, we have set *d*_max_ = 15 as a trade-off between runtime and accuracy. The best classification using this method was achieved for 13 groups with an AIC of 265 464. Using the CBN model from [27], a classification with 13 groups led to an AIC of 288 201, and the best AIC we obtained for this method was 286 496 for 50 groups.^3^ For the base-rate model, a classification with 13 groups led to an AIC of 301 506, and the best AIC we obtained was 292 902 for 8 groups.

It can clearly be seen from the AIC that allowing the model to infer pairwise influences between events provides a significant benefit, as the fastMHN model out-performs the base-rate model substantially. Additionally, the fastMHN model also achieves a significantly lower AIC than the CBN model.

While improved AIC demonstrates better statistical fit, the central question is whether this translates into biologically meaningful and clinically interpretable structure, which we investigate next.

### Independence of fastMHN classification of event count

Figure 3 shows the membership probabilities from Eq. (13) for all samples under each of the trained models. Groups obtained via the CBN and base-rate models correlate strongly with the number of active events, while groups obtained via the fastMHN model do not. A classification that correlates less strongly with *d*_active_ may be desirable in many contexts. To understand the systematic difference in classification behavior, note that both the base-rate and the CBN model calculate occurrence probabilities independently of events already present, as the base-rate model does not take interactions into account at all and the CBN model only accounts for dependencies via enforcing a partial order of events [3, 27]. In the MHN model, on the other hand, there are additional continuous parameters to describe the interaction between different events. This enables the model to describe an entire cancer-progression process for each group, and hence samples with few and many events can fit into the same group. As a result, the membership probabilities seen in Fig. 3 are more evenly distributed for the fastMHN model when very few events are present, and the model becomes more certain when the events present are distinctive enough.

This can also be seen in Fig. 4, in which we investigate to what extent membership probabilities change during the progression of a tumor. For the base-rate and CBN models, membership probabilities change more rapidly between subsequent steps of the progression. In particular the CBN model shows large confidence when only few events are present in the sample, but probabilities change drastically between steps in the progression. This behavior is expected from the strong correlation between the groups and *d*_active_ seen in Fig. 3 but can be problematic, e.g., when using the classification as a basis for therapeutic decisions, as observing the tumor at a slightly different time might lead to a drastically different course of action. The fastMHN model, on the other hand, exhibits more uniform membership probabilities in the early stages of tumor progression and only becomes more confident once more information is available. This behavior is desirable since all tumors evolve from the same starting point of healthy cells, so different dynamics can only be differentiated later in the progression. We see in Fig. 4 that membership probabilities do not change drastically between steps and the evolution of the membership probabilities is smoother overall.

**Figure 4.**
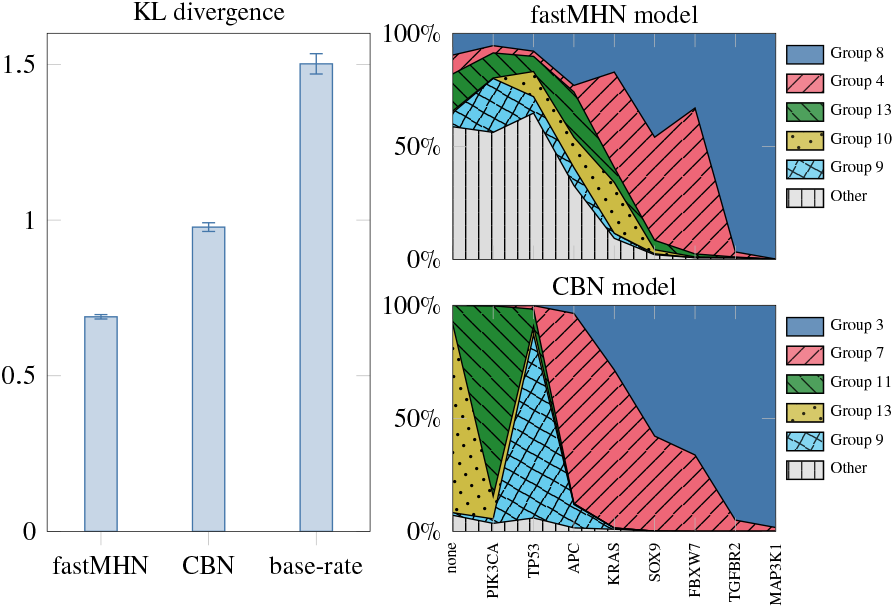
Evolution of the membership probabilities under the fastMHN, CBN, and base-rate models. Left: Average KL divergence of the membership probabilities of any observed tumor (defined by its mutational state) from the membership probabilities of any of its direct predecessors (defined as any observed tumor with exactly one mutation less). Right: Membership probabilities for a fictional tumor’s progression trajectory as it accumulates events from left (PIK3CA first) to right (MAP3K1 last). This trajectory was chosen as it is the longest chain of events that was observed in the dataset.

### Subgroup identification beyond cancer types

Next, we investigate the composition of the groups found by the fastMHN and CBN models. Figure 5 shows how many samples in each group originate from each of the five cancer types in the dataset. For both models, the composition differs significantly between groups, showing that the classification uncovers biological signals in both cases.

**Figure 5.**
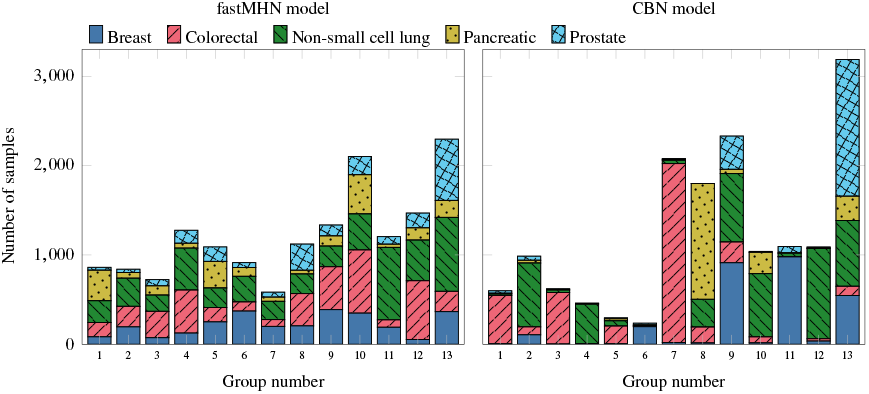
Composition of the groups found by the fastMHN and CBN models with respect to cancer types.

All fastMHN groups have significant contributions from many different cancer types, with fractions of the different types varying significantly between groups. This can be understood by looking at the fastMHN networks of the individual groups. For example, in group 11, the base rate of STK11 is the largest of any of the groups, and EGFR additionally has one of the largest base rates. Both of these events are common in non-small cell lung cancer [33, 37], so it is expected that this group is dominated by samples of this cancer type. In most other groups, such specific markers from one cancer type are absent, so group composition is more heterogeneous, as the same progression dynamics was observed for samples from different cancer types.

For the CBN model, the group composition looks very different. Here, many groups (i.e., all groups except 9 and 13) are dominated by samples from a single cancer type. This is already expected from the membership probabilities in Fig. 3, as, e.g., prostate cancers tend to have only few genetic events and will therefore not appear in many of the CBN groups, which are highly correlated with the number of active genetic events. Therefore, the CBN model tends to find subgroups within a cancer type, while the fastMHN model finds similarities in the progression across different cancer types.

We also see that group sizes are more similar for the fastMHN model, ranging from 583 samples in group 7 to 2296 samples in group 13 while for the CBN model, the sizes range from 236 to 3187 samples.

### A cross-tissue tumor subtype with poor survival linked to STK11 mutations

Finally, we investigate how the classification obtained by fastMHN reveals parallels in the progression across different cancer types and how it can be used to find de-novo subtypes. To this end we focus on group 11. Figure 5 shows that it is dominated by non-small cell lung cancer, which makes up 67 % of the samples in this group.

The samples from other cancer types in this group share similar co-occurrence patterns as the non-small lung cancer samples, as can be seen in Fig. 6. For example, both cohorts exhibit pairwise mutual exclusivity of STK11 and EGFR, and of EGFR and TP53. On the other hand, STK11 and KRAS often occur together in both cohorts. These are typical patterns for non-small lung cancer [13, 24, 49], but have not been established yet for other cancer types. A notable difference between the lung and non-lung cancers in group 11 is that the former also show a higher occurrence of KEAP1 in STK11-positive tumors. The co-occurrence of these two genetic events has been explained by their superadditive protective effects against certain cellular stressors, specifically in the lung-cancer context [48]. Furthermore, many of the non-lung cancers do not exhibit the more prevalent events. Overall, Fig. 6 shows that our classification assigns samples to group 11 because their underlying dynamics are similar.

**Figure 6.**
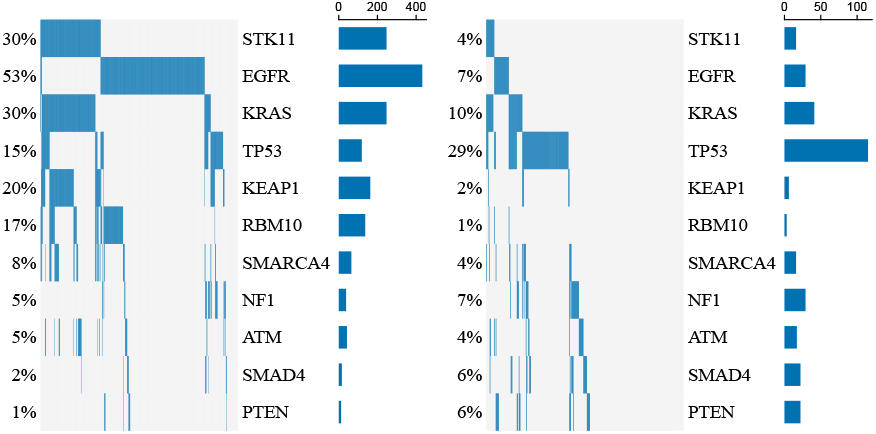
Oncoplot and histograms showing the composition of group 11 for non-small cell lung cancer (left) and other cancers (right), including only STK11 and its ten strongest interaction partners based on the sums of the absolute values of incoming and outgoing effects in the *θ* matrix of group 11. The interaction partners are sorted by overall prevalence.

In the remainder of this section, we investigate in more detail the role of STK11 in group 11 and other groups. We start with non-small cell lung cancer, for which STK11 is associated with shorter patient survival [37]. This is seen in our data as well, and the association is more pronounced in group 11 than in other groups, see the left column of Fig. 7.

**Figure 7.**
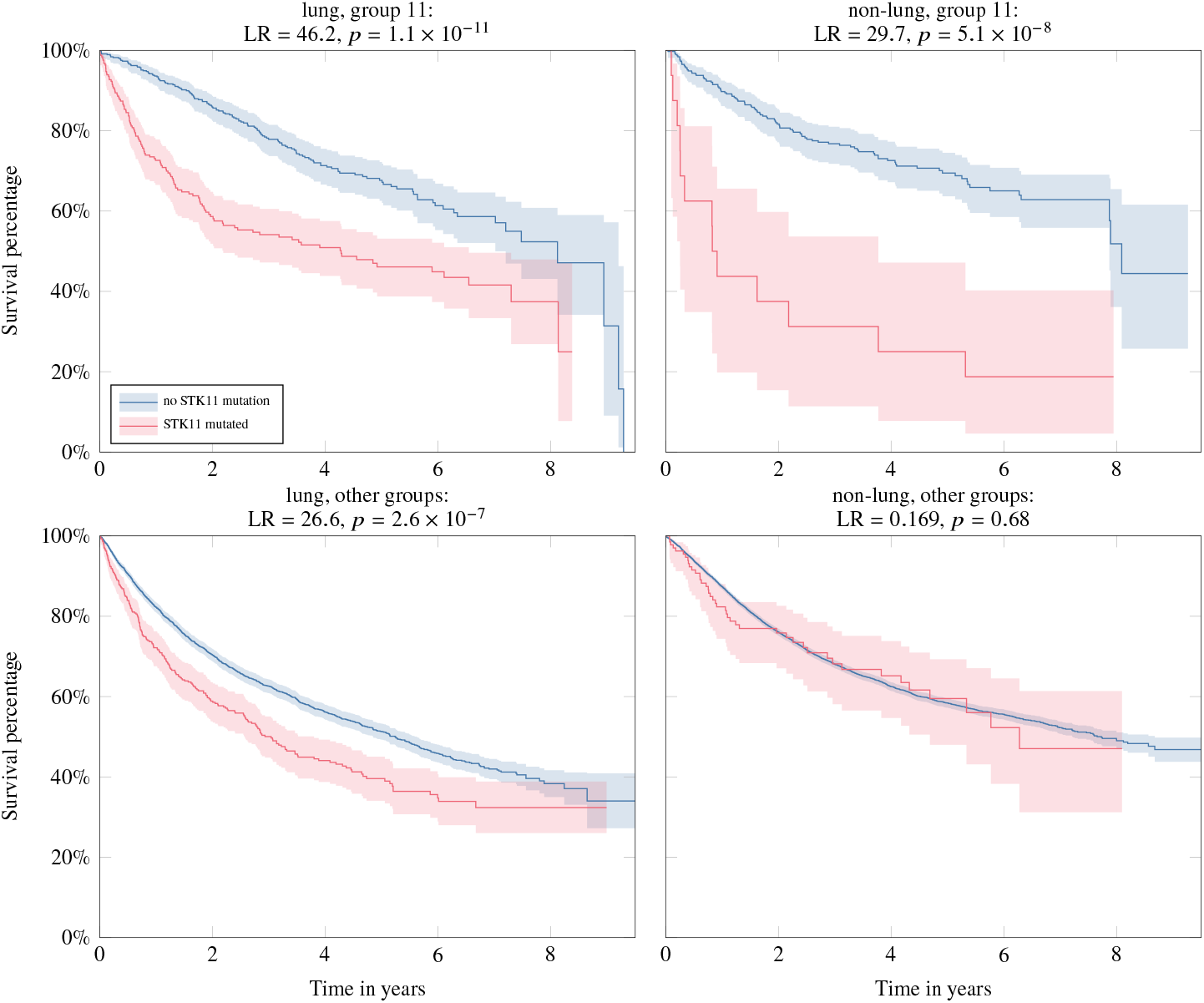
Relation between STK11 and patient survival. We compare non-small cell lung cancers (left) and other cancer types (right) for fastMHN group 11 (top) and other groups (bottom). Likelihood ratios (LR) and *p*-values of log-rank tests are shown in the individual titles.

For non-lung cancers, see right column of Fig. 7, we find that in groups other than group 11, there is no significant relation between patient survival and the presence of STK11. In group 11, however, a significant association can be seen, i.e., patients exhibiting a mutation in STK11 show much shorter survival than patients without such a mutation. 6.7 % of breast, 2.1 % of colorectal, 1.8 % of pancreatic, and 4.0 % of prostate cancer samples from our dataset are classified into group 11. These samples follow a progression that is similar to the non-small cell lung cancers in group 11, potentially enabling transfer of the knowledge of this rare molecular subtype to the non-lung cancers in group 11.

We see that our model indeed classifies samples into the same group that exhibit similar tumor-progression dynamics and show similar associations with patient survival. The example of group 11 shows that our pan-cancer classification can be used to find novel subtypes of cancer. In particular, it enables us to find groups across cancer types even when these groups are uncommon and only a limited amount of data is available for some cancer types. Using such information can help to inform therapeutic decisions in particular for cancers with atypical genetic profiles.

## DISCUSSION

In this paper we have presented fastMHN, a strategy for approximate MHN and oMHN training, breaking the exponential scaling barrier of the exact training method. We showed that this strategy leads to good approximations of the exact training process, both in each individual step as well as in the entire process.

We then applied this strategy to pan-cancer classification on a biologically relevant dataset that was out of reach of the existing oMHN training method. By comparing to a simple baseline model as well as to the CBN method [27], we showed that fastMHN gives a classification that provides a view on tumor groups that is more strongly based on co-occurence patterns governing cancer progression and that is more realistic in its group assignments.

Our results suggest that pan-cancer progression modeling can serve as a framework for transferring biological and clinical insights between oncological fields. Across tumor entities, progression mechanisms that are well characterized in one cancer type may remain largely unexplored in others, not because they are absent, but because they occur only in small molecular groups. In entity-centered analyses, such rare mechanisms are difficult to detect and are often overlooked. By aggregating evidence across tumor types, our pan-cancer analysis increases sensitivity to shared progression dynamics that would remain inaccessible in single-entity studies.

Within this framework, the groups identified by our model should not be interpreted as novel cancer entities. Rather, our results indicate that they highlight recurring cancer-progression mechanisms that manifest across tumors of different origin. The STK11-associated group provides a concrete example. While the prognostic relevance of STK11 mutations is well-established in non-small cell lung cancer, our analysis suggests that a similar context exists in molecularly defined subgroups of other cancer types, where this association has not previously been described systematically.

Importantly, in non-small cell lung cancer, STK11 mutations are known to be associated with poor response to immune checkpoint inhibitors [42] and often motivate a shift away from immunotherapy towards more intensive chemotherapy regimens. Our results suggest the clinical hypothesis that comparable considerations may apply to STK11-driven subgroups in other tumor entities. While such implications are necessarily speculative and require prospective validation, this example illustrates how pan-cancer progression modeling can move beyond descriptive classification and generate clinically relevant hypotheses. In this way, the approach has the potential not only to deepen biological understanding, but also to expand therapeutic decision-making options for patients across cancer types.

## CODE AVAILABILITY

The python package fastmhn for approximate MHN and oMHN training is available at github.com/simon-pfahler/fastmhn. The pan-cancer classification workflow and all data presented are available at github.com/simon-pfahler/fastMHN-classification.

## COMPETING INTERESTS

No competing interest is declared.

## AUTHOR CONTRIBUTIONS STATEMENT

Simon Pfahler, Tilo Wettig, and Rainer Spang conceptualized and initiated the project. Simon Pfahler, Rudolf Schill, and Tilo Wettig developed the method. Simon Pfahler implemented the algorithms. Simon Pfahler and Tilo Wettig validated the method. Simon Pfahler and Andreas Lösch prepared the input data. Simon Pfahler and Andreas Lösch analyzed the classification results. Andreas Lösch, Y. Linda Hu, Rudolf Schill, and Rainer Spang provided biological explanations of he results. Simon Pfahler, Tilo Wettig, Rainer Spang, and Y. Linda Hu drafted the manuscript. All authors critically read and improved upon the draft.

## ACKNOWLEDGEMENTS

This work is funded in part by the Free State of Bavaria through the Marianne-Plehn Programme. It is also funded in part by the German Research Foundation (DFG) through the grants “Tensorapproximationsmethoden zur Modellierung von Tumorprogression” (project 458051812) and “Striking a moving target: From mechanisms of metastatic organ colonization to novel systemic therapies” (TRR 305). It is also funded in part by the Swiss National Science Foundation through the grant “Inferring tumor phylogenies from single-cell data for dynamic patient risk prediction” (10004456).

1 Using a different mean would correspond to a simple transformation of the model parameters.

2 For example, using 20 iterations, 10 groups, and 5-fold cross-validation with typically 4 different regularization strengths leads to ~4000 trained oMHNs for one classification.

3 50 groups was the largest number of groups tested.

## Notes

### Competing Interest Statement

The authors have declared no competing interest.

## References

[1] David Barber. Bayesian Reasoning and Machine Learning. Cambridge University Press, February 2012.

[2] Fritz Bayer, Marco Roncador, Giusi Moffa, et al. Network-based clustering unveils interconnected landscapes of genomic and clinical features across myeloid malignancies. Nature Communications, 16(1):4043, April 2025.

[3] Niko Beerenwinkel, Nicholas Eriksson, and Bernd Sturmfels. Conjunctive Bayesian networks. Bernoulli, 13(4):893–909, November 2007.

[4] Peter Benner, Ren-Cang Li, and Ninoslav Truhar. On the ADI method for Sylvester equations. Journal of Computational and Applied Mathematics, 233(4):1035–1045, December 2009.

[5] Christopher M. Bishop. Pattern Recognition and Machine Learning, volume 4. Springer, 2006.

[6] Joseph E. Cavanaugh and Andrew A. Neath. The Akaike information criterion: Background, derivation, properties, application, interpretation, and refinements. WIREs Computational Statistics, 11(3):e1460, 2019.

[7] Ethan Cerami, Jianjiong Gao, Ugur Dogrusoz, et al. The cBio cancer genomics portal: An open platform for exploring multidimensional cancer genomics data. Cancer Discovery, 2(5):401–404, May 2012.

[8] Feixiong Cheng, Chuang Liu, Bairong Shen, and Zhongming Zhao. Investigating cellular network heterogeneity and modularity in cancer: A network entropy and unbalanced motif approach. BMC Systems Biology, 10(3):65, August 2016.

[9] Giovanni Ciriello, Martin L. Miller, Bülent Arman Aksoy, et al. Emerging landscape of oncogenic signatures across human cancers. Nature Genetics, 45(10):1127–1133, October 2013.

[10] David R. Cox. Regression Models and Life-Tables. Journal of the Royal Statistical Society: Series B (Methodological), 34(2):187–202, 1972.

[11] Ino de Bruijn, Ritika Kundra, Brooke Mastrogiacomo, et al. Analysis and Visualization of Longitudinal Genomic and Clinical Data from the AACR Project GENIE Biopharma Collaborative in cBioPortal. Cancer Research, 83(23):3861–3867, December 2023.

[12] Juan Diaz-Colunga and Ramon Diaz-Uriarte. Conditional prediction of consecutive tumor evolution using cancer progression models: What genotype comes next? PLOS Computational Biology, 17(12):e1009055, December 2021.

[13] Francesco Facchinetti, Maria Virginia Bluthgen, Gabrielle Tergemina-Clain, et al. LKB1/STK11 mutations in non-small cell lung cancer patients: Descriptive analysis and prognostic value. Lung Cancer, 112:62–68, October 2017.

[14] Roger Fletcher. Practical Methods of Optimization. John Wiley & Sons, 2013.

[15] K. Gallivan, A. Vandendorpe, and P. Van Dooren. Sylvester equations and projection-based model reduction. Journal of Computational and Applied Mathematics, 162(1):213–229, January 2004.

[16] Jianjiong Gao, Bülent Arman Aksoy, Ugur Dogrusoz, et al. Integrative analysis of complex cancer genomics and clinical profiles using the cBioPortal. Science Signaling, 6(269):pl1, April 2013.

[17] Peter Georg, Lars Grasedyck, Maren Klever, et al. Lowrank tensor methods for Markov chains with applications to tumor progression models. Journal of Mathematical Biology, 86(1):7, December 2022.

[18] Moritz Gerstung, Michael Baudis, Holger Moch, and Niko Beerenwinkel. Quantifying cancer progression with conjunctive Bayesian networks. Bioinformatics, 25(21):2809–2815, November 2009.

[19] Moritz Gerstung, Clemency Jolly, Ignaty Leshchiner, et al. The evolutionary history of 2,658 cancers. Nature, 578(7793):122–128, February 2020.

[20] Lars Grasedyck, Daniel Kressner, and Christine Tobler. A literature survey of low-rank tensor approximation techniques. GAMM-Mitteilungen, 36(1):53–78, 2013.

[21] Douglas Hanahan and Robert A. Weinberg. Hallmarks of cancer: The next generation. Cell, 144(5):646–674, March 2011.

[22] Katherine A. Hoadley, Christina Yau, Denise M. Wolf, et al. Multiplatform Analysis of 12 Cancer Types Reveals Molecular Classification within and across Tissues of Origin. Cell, 158(4):929–944, August 2014.

[23] Justin Jee, Christopher Fong, Karl Pichotta, et al. Automated real-world data integration improves cancer outcome prediction. Nature, 636(8043):728–736, December 2024.

[24] Ruirui Jiang, Bo Zhang, Xiaodong Teng, et al. Validating a targeted next-generation sequencing assay and profiling somatic variants in Chinese non-small cell lung cancer patients. Scientific Reports, 10(1):2070, February 2020.

[25] Hyeongmin Kim and Yong-Min Kim. Pan-cancer analysis of somatic mutations and transcriptomes reveals common functional gene clusters shared by multiple cancer types. Scientific Reports, 8(1):6041, April 2018.

[26] Diederik P. Kingma and Jimmy Ba. Adam: A Method for Stochastic Optimization. In Proceedings of the 3rd International Conference on Learning Representations (ICLR 2015), January 2017.

[27] Jack Kuipers, Thomas Thurnherr, Giusi Moffa, et al. Mutational interactions define novel cancer subgroups. Nature Communications, 9(1):4353, October 2018.

[28] Yuanyuan Li, Kai Kang, Juno M. Krahn, et al. A comprehensive genomic pan-cancer classification using The Cancer Genome Atlas gene expression data. BMC Genomics, 18(1):508, July 2017.

[29] Thomas Mach and Jens Saak. How Competitive is the ADI for Tensor Structured Equations? PAMM, 12(1):635–636, 2012.

[30] Anders L. Madsen and Finn V. Jensen. Lazy propagation: A junction tree inference algorithm based on lazy evaluation. Artificial Intelligence, 113(1):203–245, September 1999.

[31] Saeed Masoudnia and Reza Ebrahimpour. Mixture of experts: A literature survey. Artificial Intelligence Review, 42(2):275–293, August 2014.

[32] Franziska Michor, Yoh Iwasa, and Martin A. Nowak. Dynamics of cancer progression. Nature Reviews Cancer, 4(3):197–205, March 2004.

[33] Anita Midha, Simon Dearden, and Rose McCormack. EGFR mutation incidence in non-small-cell lung cancer of adenocarcinoma histology: A systematic review and global map by ethnicity (mutMapII). American Journal of Cancer Research, 5(9):2892–2911, August 2015.

[34] T.K. Moon. The expectation-maximization algorithm. IEEE Signal Processing Magazine, 13(6):47–60, November 1996.

[35] Rosanna Overholser and Ronghui Xu. Effective degrees of freedom and its application to conditional AIC for linear mixed-effects models with correlated error structures. Journal of Multivariate Analysis, 132:160–170, November 2014.

[36] Simon Pfahler, Peter Georg, Rudolf Schill, et al. Taming numerical imprecision by adapting the KL divergence to negative probabilities. Statistics and Computing, 34(5):168, August 2024.

[37] Cristiana M. Pineda, Zoe Guan, Hyunwoo Kwon, et al. A Comprehensive Mutational and Histopathological Analysis of STK11-Mutant Non-Small Cell Lung Carcinomas. Modern pathology, 39(1):100938, November 2025.

[38] Daniele Ramazzotti, Giulio Caravagna, Loes Olde Loohuis, et al. CAPRI: Efficient inference of cancer progression models from cross-sectional data. Bioinformatics, 31(18):3016–3026, September 2015.

[39] Rudolf Schill, Maren Klever, Andreas Lösch, et al. Overcoming Observation Bias for Cancer Progression Modeling. In Research in Computational Molecular Biology, pages 217–234, 2024.

[40] Rudolf Schill, Stefan Solbrig, Tilo Wettig, and Rainer Spang. Modelling cancer progression using Mutual Hazard Networks. Bioinformatics, 36(1):241–249, January 2020.

[41] Ronglai Shen, Adam B. Olshen, and Marc Ladanyi. Integrative clustering of multiple genomic data types using a joint latent variable model with application to breast and lung cancer subtype analysis. Bioinformatics, 25(22):2906–2912, November 2009.

[42] Ferdinandos Skoulidis, Michael E. Goldberg, Danielle M. Greenawalt, et al. Stk11/lkb1 mutations and pd-1 inhibitor resistance in kras-mutant lung adenocarcinoma. Cancer Discovery, 8(7):822–835, July 2018.

[43] D. C. Sorensen and A. C. Antoulas. The Sylvester equation and approximate balanced reduction. Linear Algebra and its Applications, 351–352:671–700, August 2002.

[44] Sam Thiagalingam. A cascade of modules of a network defines cancer progression. Cancer Research, 66(15):7379–7385, August 2006.

[45] Stefan Vocht, Yanren Linda Hu, Andreas Lösch, et al. mhn: A Python package for analyzing cancer progression with Mutual Hazard Networks. Bioinformatics Advances, 6(1):vbaf283, January 2026.

[46] Jianlin Wang, Jiao Zhang, Xuebing Dai, et al. Computational models for pan-cancer classification based on multi-omics data. Frontiers in Genetics, 16, October 2025.

[47] Robert A. Weinberg. The Biology of Cancer. W.W. Norton & Company, New York, June 2006.

[48] Corrin A. Wohlhieter, Allison L. Richards, Fathema Uddin, et al. Concurrent mutations in stk11 and keap1 promote ferroptosis protection and scd1 dependence in lung cancer. Cell Reports, 33(9):108444, December 2020.

[49] Dandan Yin, Xiyi Lu, Xiao Liang, et al. STK11 genetic alterations in metastatic EGFR mutant lung cancer. Scientific Reports, 15(1):5729, February 2025.

[50] Wei Yuan, Yijiang Chen, Biyue Zhu, et al. Pancancer outcome prediction via a unified weakly supervised deep learning model. Signal Transduction and Targeted Therapy, 10(1):285, September 2025.

